# Computational estimation of ms-sec atomistic folding times

**DOI:** 10.1101/427393

**Authors:** Upendra Adhikari, Barmak Mostofian, Jeremy Copperman, Andrew Petersen, Daniel M. Zuckerman

## Abstract

Despite the development of massively parallel computing hardware including inexpensive graphics processing units (GPUs), it has remained infeasible to simulate the folding of atomistic proteins at room temperature using conventional molecular dynamics (MD) beyond the µs scale. Here we report the folding of atomistic, implicitly solvated protein systems with folding times τ_f_ ranging from ∼100 µs to ∼1s using the weighted ensemble (WE) strategy in combination with GPU computing. Starting from an initial structure or set of structures, WE organizes an ensemble of GPU-accelerated MD trajectory segments via intermittent pruning and replication events to generate statistically unbiased estimates of rate constants for rare events such as folding; no biasing forces are used. Although the variance among atomistic WE folding runs is significant, multiple independent runs are used to reduce and quantify statistical uncertainty. Folding times are estimated directly from WE probability flux and from history-augmented Markov analysis of the WE data. Three systems were examined: NTL9 at low solvent viscosity (yielding τ_f_ = 0.8 − 9.0 *μs*), NTL9 at water-like viscosity (τ_f_ = 0.2 − 1.9 ms), and Protein G at low viscosity (τ_f_ = 3.3 - 200 ms). In all cases the folding time, uncertainty, and ensemble properties could be estimated from WE simulation; for Protein G, this characterization required significantly less overall computing than would be required to observe a *single folding event* with conventional MD simulations. Our results suggest that the use and calibration of force fields and solvent models for precise estimation of *kinetic* quantities is becoming feasible.

## Introduction

Elucidating the kinetics and mechanisms of protein folding has been a decades-long focus of molecular biophysics, both experimental and theoretical/computational.^1-19^ Significant challenges remain, however, notably whether molecular dynamics (MD) simulations will provide the hoped-for reproducible and atomically detailed folding trajectories.^1, 11, 13-14, 20-23^ Despite isolated reports of success,^24-25^ MD simulations generally have not produced *room temperature* atomistic folding trajectories beyond the µs timescale even with modern hardware.^26^ Promising results have been reported using path-sampling techniques^27-31^ but no simulation methodology has emerged as a general-purpose tool for folding, especially for timescales beyond the µs range.

Here we report substantial progress in the application of the weighted ensemble (WE) path sampling method^32-36^ to room-temperature folding at the microsecond (μs), millisecond (ms) and second (s) scales, exploiting the power of GPU and cluster computing. We study three atomistic implicitly solvated systems: NTL9 with low and high-friction solvent, as well as Protein G at low friction. These are costly studies, requiring aggregate trajectory totals of 10s to 100s of µs per system, but they enable fairly precise (order-of-magnitude) estimation of folding rate constants. In earlier work, Ensign and Pande^26^ were able to estimate the WW-domain folding time of ∼100 µs at room temperature using distributed computing with a total cost of 400 – 500 µs per system. To our knowledge, there are no other computations of room-temperature atomistic protein folding rates at the ms scale and beyond. Prior folding-rate calculations of NTL9 and Protein G were conducted at high temperature (355 K^1^/370 K^37^, and 350 K^1^ respectively) because of the prohibitive room-temperature timescales.

In addition to information about protein folding, the ability to quantify rate constants for slow-timescale biomolecular behavior is a critical step in model (force field) development. Although MD simulation is now a standard tool in structural biology studies,^38-41^ the governing parameters of MD force fields have been determined based on energy minima^42-46^ whereas energy barriers are expected to govern kinetic behavior. Given the evident importance of dynamic biomolecular phenomena, it is critical to obtain simulation-based rate constants to permit further refinement of force fields. Force fields cannot be assessed fully without the ability to compute kinetic observables, and we report on significant progress in this regard.

The WE method (Fig. 1A) employed in the present report is one of a number of path sampling approaches based on rigorous statistical mechanics^33, 47-52^ capable of yielding unbiased rate constants. Although all these methods are theoretically well-grounded, WE does offer the pragmatic advantage of being fully independent of the dynamics engine employed, which has enabled its application with a wide range of both molecular and cell-scale simulation software.^34, 53-59^ This versatility facilitated the integration of the WESTPA software package^60^ with the GPU-accelerated version of the AMBER molecular dynamics package^61-63^ as employed here. The WE method yields ensembles of fully continuous trajectories from which non-equilibrium observables can be calculated, including kinetic and mechanistic properties. Importantly, the continuous WE trajectories spanning from unfolded to folded macrostates enable folding rate estimation using the history-augmented Markov state model (haMSM) formalism which is unbiased at arbitrary lag times.^56, 64^

**Figure 1:**
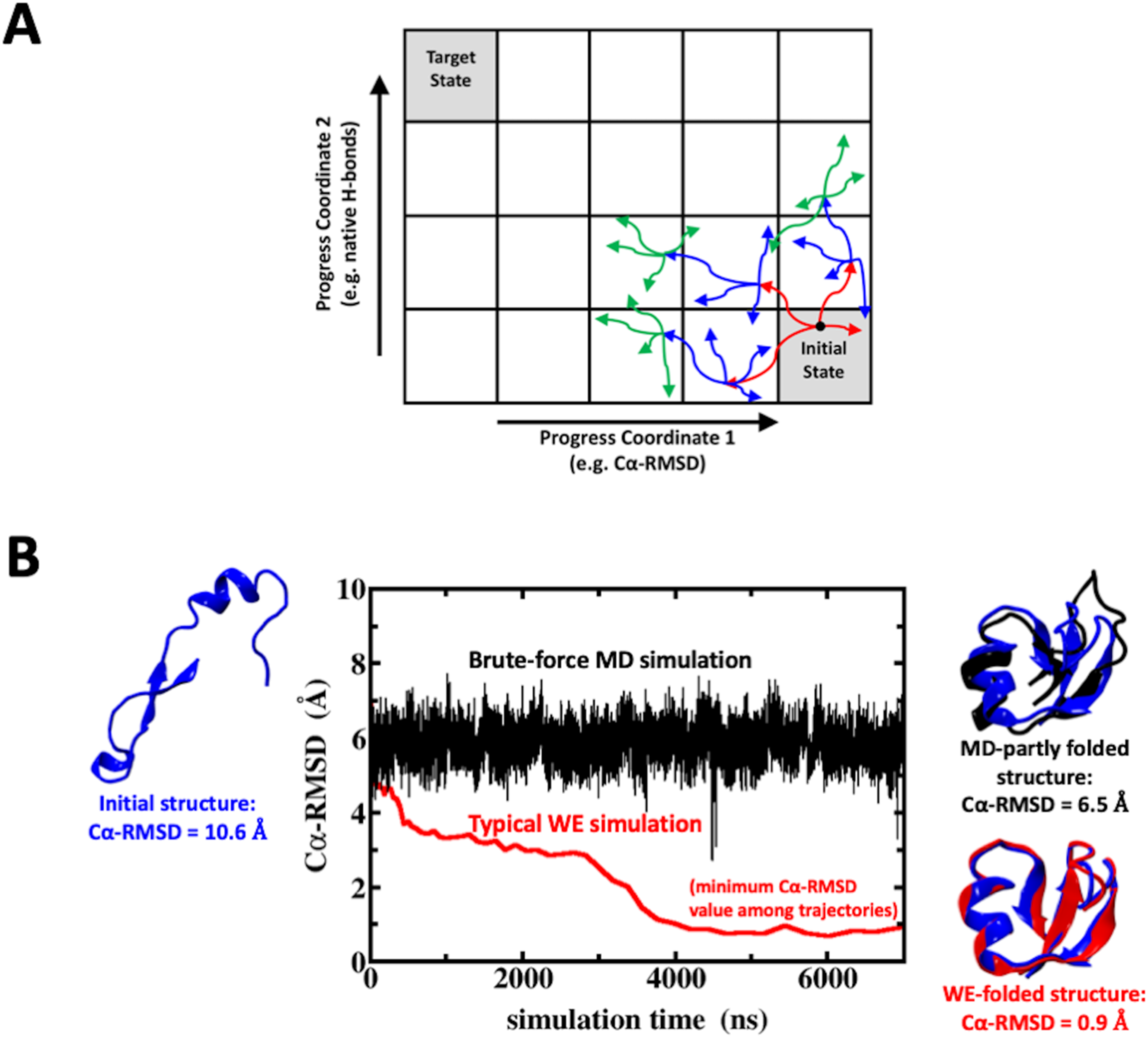
The WE procedure and comparison to regular MD simulation. (A) A schematic of the WE simulation procedure is shown with two-dimensional binning for protein folding. Three iterations (red, then blue, then green) are shown based on a target number of 4 trajectories per bin, illustrating the “statistical ratcheting” effect which is possible without applying biasing forces. Note that a set of new trajectories is shown only for those parent trajectories that reached a new bin. (B) A brute-force MD simulation of NTL9 leads to the “MD-partly folded” structure (black structure) with a Cα-RMSD of 6.5 Å with respect to the folded crystal structure (blue structures at right) after 7 μs of simulation time. By contrast, a WE simulation starting from the same initial structure (blue structure at left) samples the “WE-folded” NTL9 structure with Cα-RMSD < 1 Å (red structure on the right panel). The WE simulation time is the aggregate time including all trajectory segments, representing a fair comparison using roughly the same amount of computational resources.

## Results

The WE procedure takes advantage of running in parallel multiple simulations with well-defined probabilities (or weights) in a conformational space that typically is divided based on pre-defined progress coordinates (see Fig. 1A).^32^ The trajectory pruning and replication strategy facilitates progress along the coordinates and guarantees a constant total weight of all trajectories during the WE simulation (see SI Methods for more details). Fig. 1B shows a comparison of a brute-force MD simulation with a typical WE simulation, both starting from the same unfolded NTL9 structure. After ∼7 μs of aggregate simulation time, the NTL9 Cα-RMSD in the MD simulation remains > 6 Å, whereas in the WE simulation folded NTL9 structures with Cα-RMSD < 1 Å are sampled. The probability flux of simulations reaching the target state allows estimation of the folding kinetics and the interrogation of continuous trajectories can provide information on folding mechanisms.

Estimates for folding rate constants are derived in two ways from WE data – directly from observed probability flows and using haMSM analysis. In both approaches, all WE simulations were used for a given system. In the direct analysis, Figs. 2-4 show that the probability flux into the folded states, which is an estimator for the rate constant,^65^ apparently reaches a steady value in all three atomistic folding systems: NTL9 at low friction, NTL9 at high friction, and Protein G at low friction. The “molecular time”, t_mol_, shown in Figs. 2-4 represents the time elapsed during individual trajectories. It is noteworthy that the flux reaches a plateau rather abruptly for Protein G, and to a value lower than the experimental value despite the low friction model, in contrast to the NTL9 data. Although the folding flux is dominated by a relatively small fraction of the independent runs, the dominating runs switch during the course of the trajectories (Figs. S1-S3). Nevertheless, the profiles of flux vs. Cα-RMSD (Figs. S4-S6) indicate the Protein G simulations in fact are far from steady state, implying the direct flux value for that system is not reliable despite its apparent plateau as a function of t_mol_. The flux profiles at true steady state should be constant at all hypersurfaces (e.g., fixed Cα-RMSD values) separating folded from unfolded macrostates.^66-67^

**Figure 2:**
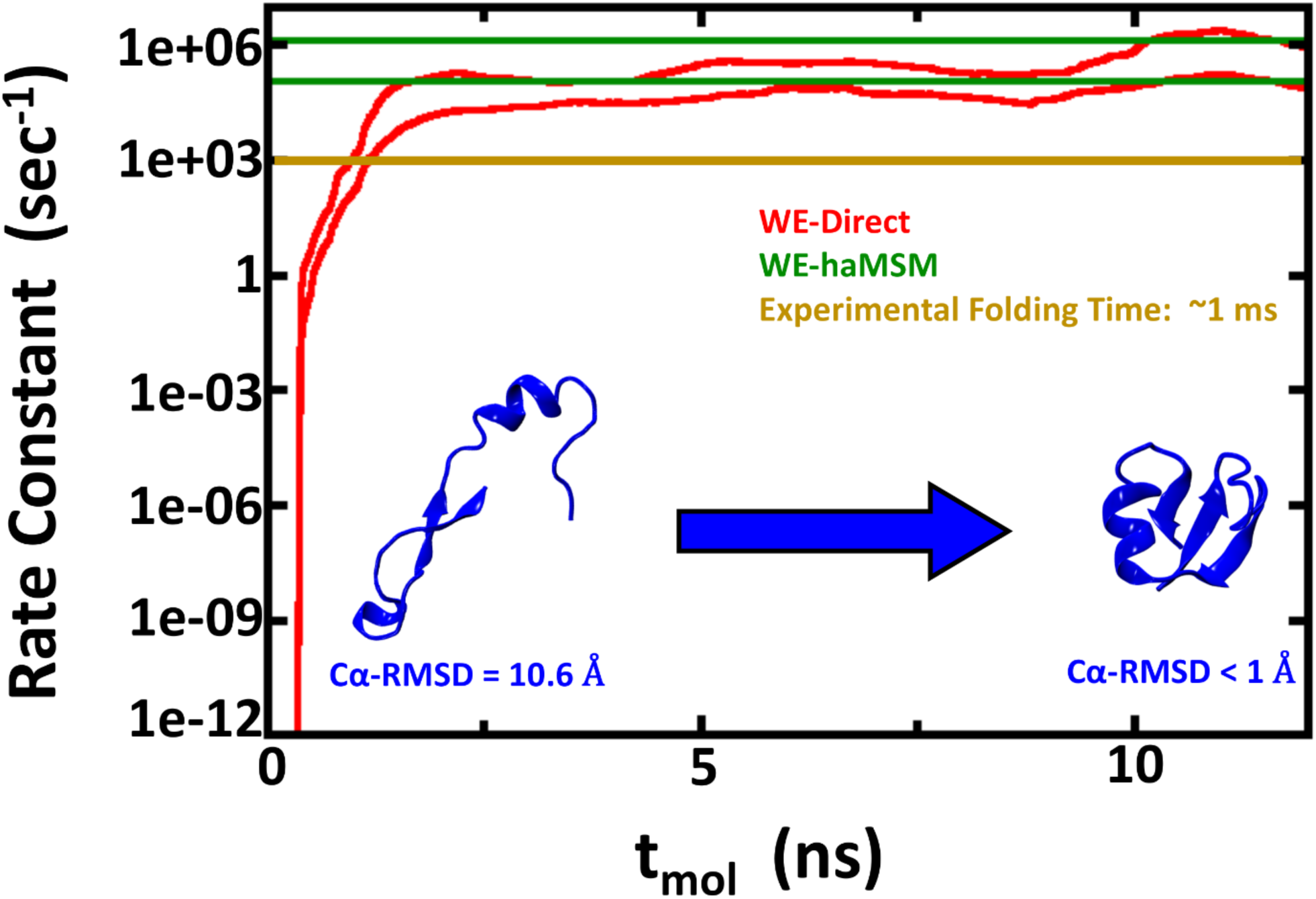
Rate constant estimations for NTL9 folding using 2D WE method with solvent viscosity (γ) set to 5 ps^−1^. The red lines show the nominal 95% Credibility Region (CR) as a function of molecular time from Bayesian bootstrapping based on direct WE rate constant estimates, which were windowed averages of the previous 1 ns of molecular time for each of the 10 independent simulations (see Figure S1). The green lines show the 95% CR for rate constants obtained by the haMSM method. The experimental rate constant is shown in gold, but note that the low viscosity used in these simulations is expected to yield overly fast kinetics.

**Figure 3:**
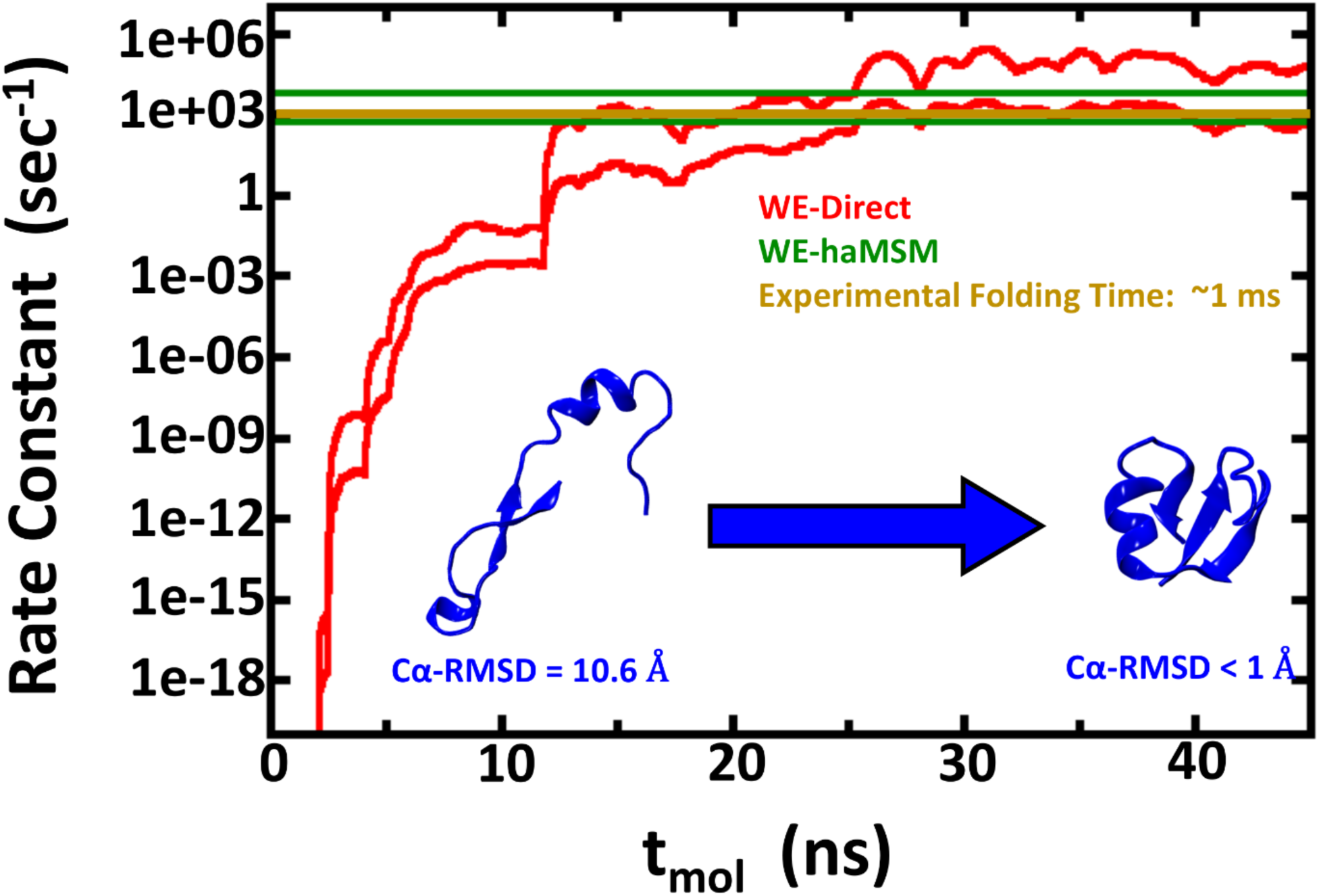
Rate constant estimations for NTL9 folding using 1D WE method with solvent viscosity (γ) set to 80 ps^−1^. The red lines show the nominal 95% Credibility Region (CR) as a function of molecular time from Bayesian bootstrapping based on direct WE rate constant estimates, which were windowed averages of the previous 1 ns of molecular time for each of the 30 independent simulations (see Figure S2). The green lines show the 95% CR for rate constants obtained by the haMSM method. The experimental rate constant is shown in gold.

**Figure 4:**
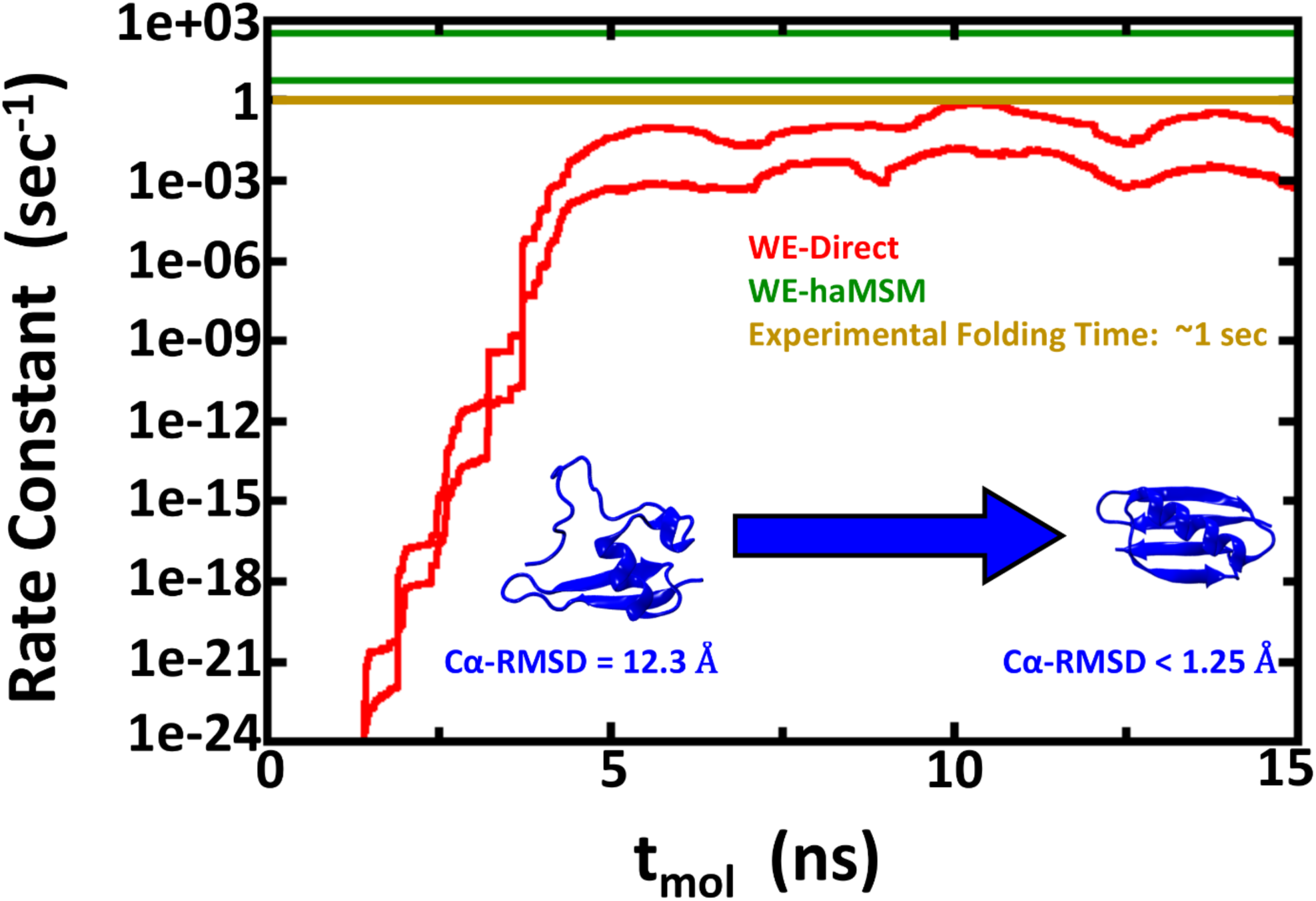
Rate constant estimations for Protein G folding using 2D WE method with solvent viscosity (γ) set to 5 ps^−1^. The red lines show the nominal 95% Credibility Region (CR) as a function of molecular time from Bayesian bootstrapping based on direct WE rate constant estimates, which were windowed averages of the previous 1 ns of molecular time for each of the 15 independent simulations (see Figure S3). The green lines show the 95% CR for rate constants obtained by the haMSM method. The experimental rate constant is shown in gold, but note that the low viscosity used in these simulations is expected to yield overly fast kinetics.

We also employed haMSM analysis, which is unbiased for steady-state flux estimation at arbitrary lag times, and small lag times allow fuller use of the extensive WE data.^56, 64, 68^ The approach is of particular interest for Protein G because, in principle, a haMSM can estimate steady-state behavior using trajectories generated in the transient period – i.e., in the approach to steady state. As noted, the flux profile for Protein G indicates those WE simulations clearly remained in the transient regime. The haMSM results are also shown in Figs. 2-4. For the NTL9 systems, the haMSM rate estimates are consistent with estimates based on direct WE fluxes. For Protein G, the haMSM folding rate estimate is substantially higher than the direct WE estimate, and notably, it slightly exceeds the experimental value as expected for the low-friction solvent model.

WE simulation uses an ensemble of trajectories which all require computing resources, and aggregate simulation times are given in Table 1 (see SI for WE parameters and computing resources). Additional runs for the NTL9 systems were performed with alternative WE protocols to confirm the consistent, unbiased nature of the data: Figs. S7 and S8 show consistent time evolution of the folding flux based on different WE protocols for both low and high-friction systems.

**Table 1:** Computational cost and folding rate constants for the three systems studied here (AMBER FF14SB force field).

| System | Number of indep. WE simulations | Molec. time (ns) | Aggregate simulation time (μs) | Wall-clock time* (days /simulation) | WE-Direct folding time** | WE-haMSM folding time*** | Expt'l folding time |
| --- | --- | --- | --- | --- | --- | --- | --- |
| NTL9, $\gamma = 5 \text{ ps}^{-1}$ | 10 | 12 | 115 | 22 | 0.6 – 8.4 μs | 0.8 – 9.0 μs | ~ 1 ms |
| NTL9, $\gamma = 80 \text{ ps}^{-1}$ | 30 | 45 | 252 | 20 | 0.01 – 0.80 ms | 0.2 – 1.9 ms | ~ 1 ms |
| Protein G, $\gamma = 5 \text{ ps}^{-1}$ | 15 | 15 | 225 | 31 | 4 – 210 sec | 3.3 – 200 ms | ~ 1 sec |

\* Based on 1 GPU card or ~48 CPU cores.
\*\*Averaged from molecular times 10-12 ns (NTL9, $\gamma = 5 \text{ ps}^{-1}$ ), 30-45 ns (NTL9, $\gamma = 80 \text{ ps}^{-1}$ ), and 10-15 ns (Protein G), respectively. Given range is the nominal 95% Bayesian Credibility Region, which corresponds to a ~60% range of uncertainty: see Ref. 92 and SI.
\*\*\* From haMSMs with 10,000 microstates trained from molecular time 10 -12 ns (NTL9, $\gamma = 5 \text{ ps}^{-1}$ ), 30-45 ns (NTL9, $\gamma = 80 \text{ ps}^{-1}$ ), and 10-15 ns (Protein G), respectively. The uncertainty range is the nominal 95% Bayesian Credibility Region.

The present study necessarily estimated folding times specific to the chosen force field and solvent model, and also conditioned on the starting structures. The novelty of the results is their relatively high precision and unbiased nature due to the theoretical foundations of the WE and haMSM methods.^35, 56, 64^ Hence, although comparison to experimental folding times are shown in Table 1, readers are cautioned that the present study should be considered a first step in assessment of molecular models and initial ensembles. Given these caveats, the rough agreement with experimental values is encouraging but also points to the need for further investigation of solvent modeling and initial ensembles as discussed below.

A comparison of the force field-specific folding times and the aggregate simulation times as given in Table 1 enables assessment of the effectiveness of the WE protocol. In the case where WE exhibits least enhancement of sampling, namely NTL9 at low friction (Fig. 2), the calculated folding time range of 0.8 – 9.0 μs from Bayesian bootstrapping of haMSM estimates employed ∼100μs of aggregate simulation. Fig. 2 reveals that much of the computation was used to confirm steady behavior and in fact the folding time could have been inferred from substantially less computation. In principle, similar results could have been obtained via 5-10 independent standard MD runs totaling the same aggregate simulation time. However, given the experimental ms folding time, it is unlikely such MD runs would have been attempted, and WE provided a reliable estimate in an affordable amount of computing effort. The higher-friction NTL9 study, which should be a better mimic of aqueous viscosity,^69-71^ reveals a WE-haMSM folding time range of 0.2 – 1.9 ms (Fig. 3) that is essentially prohibitive for harvesting multiple events via conventional MD, even on modern GPU platforms. The value of the WE protocol is unambiguous for the slower Protein G system, where a folding time range of 3.3 – 200 ms is estimated in much less than a ms of aggregate simulation time (Fig. 4). By comparison, the computational cost of rate estimation here is substantially less than the previously reported overall cost of ∼500 µs to estimate a ∼65 µs room-temperature folding time.^26^

During the WE process, a variety of folding trajectories are simulated, enabling unbiased computation of ensemble properties. The weighted distributions of Cα-RMSD values shown in Figs. S9A, S10A for the NTL9 simulations and in Fig. S11A for the Protein G simulation serve as effective folding free energy profiles, which indicate that NTL9 folding has an energy minimum at Cα-RMSD = ∼6 Å and Protein G at Cα-RMSD = ∼10 Å. These regions are separated from the folded state by a free energy barrier, suggesting a definition of the transition region and thus allowing calculation of the transition times (event durations) of the continuous WE folding trajectories. Of growing interest,^72-73^ the event duration depends on the exact event starting point and on the solvent viscosity.^74-75^ For NTL9, at low viscosity, the distributions of event duration have a peak at 1.5 - 2 ns (Fig. S9B), while at the higher water-like viscosity the peaks occur at slightly larger values ∼4-5 ns (Fig. S10B). For Protein G, the event duration peaks are less clearly defined but occur in the range of ∼2-7 ns (Fig. S11B).

A visual analysis of representative intermediate structures sheds light on the folding mechanisms. The NTL9 molecular structures shown in Fig. 5A illustrate that during the folding process the α-helix is formed first, followed by the formation of the N-terminal β-hairpin. A putative rate-limiting step of NTL9 folding is characterized by the association of the C-terminal β-strand with the N-terminal β-hairpin through hydrogen bonds. During the final steps (1 Å < Cα-RMSD < 4 Å), the protein reduces its solvent-accessible surface area by ∼5 nm^2^ when forming the remaining native hydrogen bonds, bending the N-terminal β-hairpin turn, and aligning the α-helix with the β-sheet. Similarly, Protein G (Fig. 5B) folds by first forming the α-helix and both β-hairpins and then bringing them all closer to each other, which appears to define the main free energy barrier, before connecting the two hairpins with hydrogen bonds and establishing the 4-stranded β-sheet. From the initial formation of the secondary structural elements to the fully folded structure (i.e. 1 Å < Cα-RMSD < 10 Å), Protein G reduces its overall surface area by ∼ 8nm^2^.

**Figure 5:**
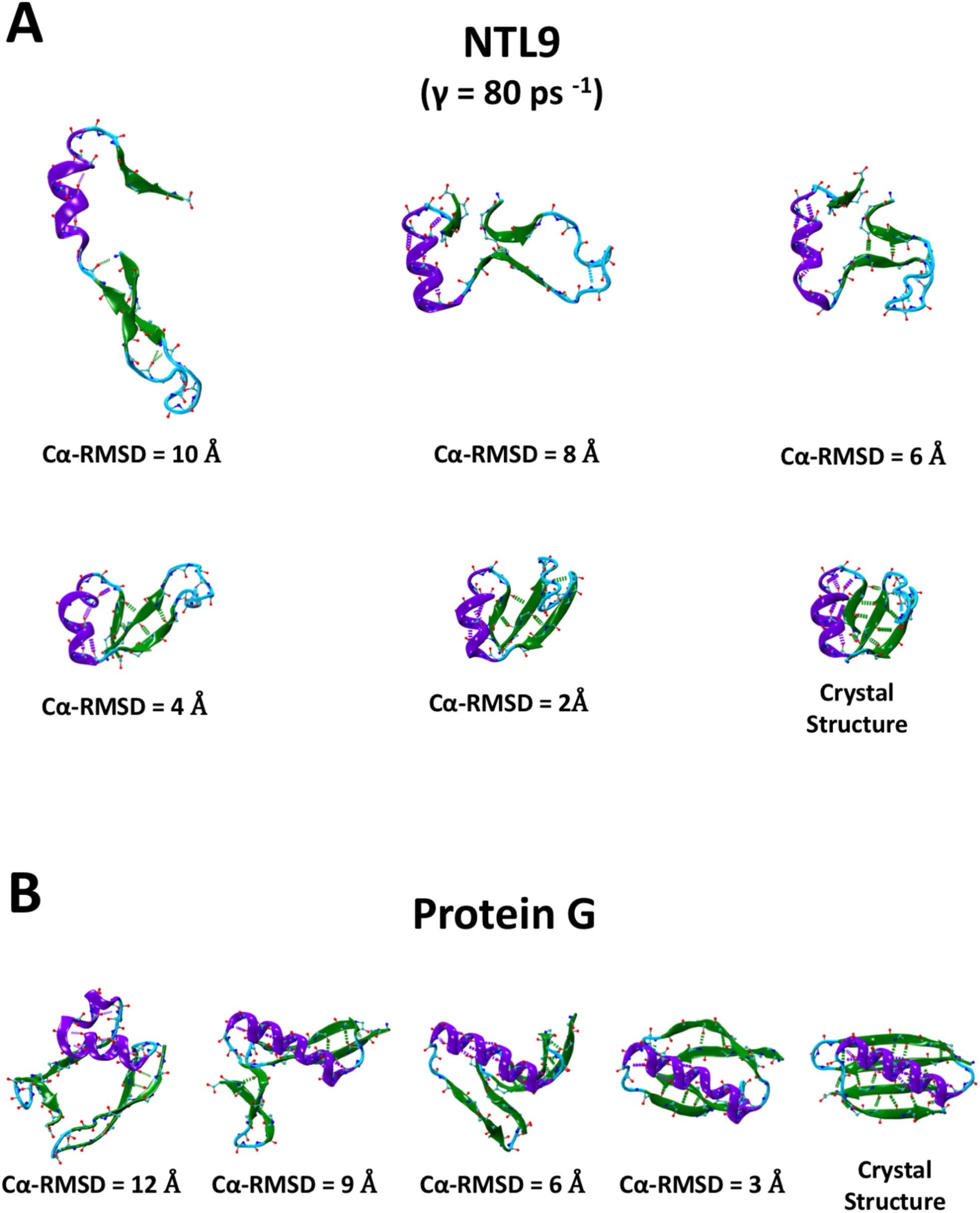
A set of example NTL9 (A) and Protein G (B) structures with decreasing Cα-RMSDs from left to right obtained from a continuous trajectory along with the folded crystal structure. Residues are colored based on their native secondary structures in violet (α-helix), green (β-sheet), and cyan (loops). Native backbone hydrogen bonds are indicated as dashed lines, if they emerge in the structure shown.

Because some prior folding studies have been performed near the melting temperature, Tm, to improve sampling,^1, 25^ it is of interest to investigate the effects of temperature on the folding process. After melting temperatures were estimated approximately (Figs. S12), we performed an additional set of WE simulations for low-friction NTL9 at Tm ∼ 325K. Comparison of the two simulation sets shows similar folding kinetics (Fig. S13) which we emphasize were obtained in the context of a single starting structure and implicit solvent. In future work, it will be of interest to compare folding mechanisms when the folding process is modeled more completely and accurately.

## Discussion

The data reported here suggest that molecular dynamics calculations may soon be able to measure precisely and regularly a broad array of experimentally relevant timescales characterizing functional motions of biomolecules. Such measurements are necessarily limited by the accuracy of the underlying model equations (i.e., the force field) but understanding and correcting force field mis-calibrations is essential for progress in computational structural biology. These corrections will not be possible without reliable kinetics measurements, and the present data yields roughly order-of-magnitude precision (Table 1). Current force fields can suffer inaccuracies exceeding 1 kcal/mol for free energy *minima*^76-78^ and errors at least as large are expected for the barriers which govern kinetics, which have not been part of force field parametrization.^21, 42-43, 79-82^ Note for reference that an order-of-magnitude change in an Arrhenius factor exp(−ΔG/RT) corresponds to a shift in Δ*G* of 1.4 kcal/mol; hence uncertainty of only 0.7 kcal/mol corresponds to a tenfold range.

Accuracy in kinetics also depends on the solvent model. Implicit solvation was employed in the present study, i.e., water molecules were not explicitly modeled. Because such models are in common use,^25, 37, 83-86^ it is important to assess their kinetic accuracy. Although the overall computational cost for WE-based rate estimation is higher at water-like viscosity (γ=80 ps^−1^), the estimated average rate constants are found to be only slightly higher than at low solvent viscosity (γ=5 ps^−1^), consistent with prior investigation of the issue.^26^ Going forward, additional comparison to explicit-solvent folding rate constants will be an important goal.

Another limitation of the present study is also intrinsic to protein folding generally – namely, ambiguity regarding the unfolded state ensemble. Experimentally, proteins are denatured chemically or with temperature,^87-90^ each of which should yield a different unfolded ensemble, and the sensitivity of refolding to the denaturing process is an under-explored topic.^91^ Given that some folding times are ms-scale or less, measurements may be sensitive to experimental protocols (e.g., mixing, cooling) occurring on the same timescales. Because of these ambiguities, we chose to keep our study as controlled as possible and focused specifically on folding from a *single* initial structure, recognizing the importance of future study of ensemble-initialized folding. Our mechanistic discussion above must be seen as restricted to this condition.

Quantification of statistical uncertainty was a central part of this study, and numerous repeated WE simulations were required to overcome the large variance of the present folding protocol (see Figs. S1-S3). Although a large variance is generally and rightly a cause for concern in data analysis, our ability to perform tens of truly independent simulations distinguishes this work from typical molecular simulation studies. As described elsewhere, neither traditional standard-error analysis nor bootstrapping properly quantify uncertainty in small-size data sets with large log-variance.^92^ We therefore employed a Bayesian bootstrapping approach both for direct WE and haMSM flux estimates, which is superior at characterizing precision in such data.^92-93^ Nevertheless, no analysis method can correct for insufficient sampling of an unknown distribution, and we estimate that the nominal 95% Bayesian credibility regions reported here empirically correspond to ∼60% probability of bracketing the true mean – and such uncertainty in the error analysis is intrinsic to the modest sample sizes.^92^ This point is borne out by the apparent ‘false plateau’ of the Protein G direct flux. Future studies will clearly benefit from variance-reduction strategies, which have been proposed.^94-95^

The weighted ensemble method was chosen over other rigorous path sampling approaches^10, 27-31, 47-52^ and standard (history-independent) Markov state models (MSMs).^96-97^ Compared to other path sampling methods, WE offers fully scalable parallelization and does not require hard-coding within the dynamics engine in order to “catch” trajectories as they cross interfaces.^34^ When compared to standard MSMs, WE not only avoids any approximation but also offers continuous trajectories and the fine temporal resolution needed to infer mechanistic details occurring on 5- 10 ns timescales (Figs. S9-S11). By contrast, modern well-validated MSMs often require lag times >100 ns.^96-97^ The continuous trajectories generated by WE allow application of the history-augmented MSMs at arbitrary lag times, which are unbiased for estimation of steady-state fluxes.^56, 64^ Using short lag-times for haMSMs in turn allows use of all data generated.

## Acknowledgements

We gratefully acknowledge support from the NIH (Grant GM115805) and from the OHSU Center for Spatial Systems Biomedicine. Computing support was provided by the Center for Research Computing at the University of Pittsburgh. Helpful comments on the manuscript were provided by Lillian Chong.

