## Supporting Information for "Computational estimation of ms-sec atomistic folding times"

### Computational estimation of ms-sec atomistic folding rates
Supporting Information

Upendra Adhikari†, Barmak Mostofian†, Jeremy Copperman†, Andrew Petersen‡, and Daniel M. Zuckerman†*
*†Department of Biomedical Engineering, School of Medicine, Oregon Health & Science University, Portland, OR 97219*
*‡ NCSU Data Science Resources, North Carolina State University, Raleigh, NC 27695*

*

#### Methods

**Systems and Molecular Dynamics simulations:**

Folded structures were taken from the Protein Data Bank crystal structures (NTL9 (residues 1 – 39) PDB ID: 2HBB,[1] Protein G (residues 1 - 56) PDB ID: 1PGA[2]). Almost all molecular dynamics simulations were performed using the AMBER 16[3-4] software package with their custom-designed hybrid precision model[5] on NVIDIA GPUs (Titan-X and GTX-1080) with a single GPU per WESTPA simulation; some NTL9 simulations with γ = 80 $ps^{-1}$ were run on CPUs. The AMBER force field FF14SB[6] was used in conjunction with Hawkins, Cramer, Truhlar[7-8] pairwise Generalized Born (GB, igb=1) implicit solvation model. A single 2 μs MD simulation of folded structures was performed to test the stability of NTL9 and Protein G structures in this force field. We also tested other solvent model combinations such as FF14SB (igb=2 & 5) and FF99SB (igb = 1, 2 & 5), which were found not suitable for these protein folding simulations as shown by a quick unfolding of folded structures with these force fields. All simulations were carried out at 300 K temperature, using a leapfrog stochastic dynamics integrator. The integration time step was taken to be 2 fs. The temperature was controlled using Langevin dynamics, and hydrogen bond length was constrained using the SHAKE algorithm. Simulations were performed at two different solvent viscosities (γ): a low solvent viscosity (γ = 5 $ps^{-1}$) and water-like viscosity (γ = 80 $ps^{-1}$) to test its effect on rate estimation.

**Starting structures for folding simulations:**

To generate starting structures, we ran unfolding simulations of both NTL9 and Protein G crystal structures using the WE method to obtain an ensemble of unfolded structures. The same WE parameters were used for both folding and unfolding simulations. Conformations with Cα-RMSD equal to or greater than 10 Å (for NTL9) and 12 Å (for Protein G) were considered to be unfolded, based on the data in the literature.[9-10] We randomly picked an unfolded structure with Cα-RMSD ~ 10.0 Å (NTL9), and ~ 12.0 Å (Protein G) from an ensemble of WE-unfolded structures to start our folding simulations. There can be many unfolded configurations which presents a difficulty on how to select an ensemble of unfolded structure. The unfolded structures selected here are not representative of all unfolded configurations. Rather, the results presented here are to show the effectiveness of the WE simulation to study the folding kinetics of both fast and slow folding systems. This work is just a beginning of the study of the estimation of folding rate constants using WE simulations. In the future, one can either take several starting structures or run independent WE simulations on them or consider starting a WE simulation with an ensemble of starting structures.

**Weighted Ensemble simulation**

Weighted ensemble simulations were carried out using the WESTPA software package.[11] In this method, multiple simulations run in parallel in a conformational space systematically divided into bins, based on some progress coordinates. Each bin may contain a fixed number (M) of trajectories or "walkers" which carry a certain weight. The simulations are periodically stopped, and the trajectories are replicated if there are less than M number of trajectories in a bin, whereas they are merged if there are greater than M number of trajectories. The total weight of the walkers is constant throughout the simulations; trajectory replication and merging are carried using statistical resampling, making the method statistically unbiased. Detailed description about WE method and WESTPA software can be found elsewhere.[11-13]

We ran non-equilibrium steady-state WE simulations for all folding data reported, where the unfolded and folded states are the initial and target states, respectively. Details regarding state definitions are given below. In non-equilibrium steady state simulations, once trajectories reach the target state, their weight is re-assigned to the initial state. The flux of the trajectories reaching the target state allows the estimation of the rate constant *k* via the Hill relation,[14-15]

$$k = \frac{1}{\text{MFPT (Unfolded}\rightarrow\text{Folded)}} = \text{Flux (Unfolded}\rightarrow\text{Folded)}$$

which over time will reach a steady value.

*WE Parameters:* The folded target states are defined to have a Cα-RMSD of 1 Å (NTL9) and 1.25 Å (Protein G), respectively, based on the brute force simulations of the folded structures. After exploring a variety of WE parameters, a resampling time (τ) of 10 ps, and four trajectories per bin were chosen. For NTL9, we ran both one dimensional (1D) and two dimensional (2D) WE simulations combined with low viscosity ($\gamma$=5 ps$^{-1}$), and water-like viscosity ($\gamma$=80 ps$^{-1}$). For Protein G, only 2D simulations with $\gamma$=5 ps$^{-1}$ were performed. In 1D simulations, Cα-RMSD was taken as the progress coordinate, whereas for 2D WE simulations, the number of H-bonds were considered as a second progress coordinate in addition to Cα-RMSD. In particular, for NTL9 we used as a second coordinate the number of H-bonds formed between any native donor and acceptor site; for protein G, we used the number of native H-bonds formed only in the alpha-helix region, as this was found to work effectively. Below, we refer to these simply as 'native H-bonds' for both systems.

Except for Figures S7 and S8, in which different WE protocols are compared for the NTL9 simulations, all results on NTL9 reported in the main text and in this supplement are based on the 2D WE simulation with $\gamma$=5 ps$^{-1}$ and the 1D WE simulation with $\gamma$=80 ps$^{-1}$. For all sets of folding simulations, multiple independent WE folding simulations were performed starting from the same initial unfolded structure and with the same simulation parameters. Rate constants were estimated for each independent run by averaging over the last 100 iterations (1 ns) to reduce the noise. A summary of WE parameters, the bin mapping, and the computational resources used in this study is given below:

Table S1: WE Parameters

| | |
|---|---|
| Resampling time (τ) | 10 ps |
| Trajectories per bin | 4 |
| Progress coordinates | Cα-RMSD (1D), + Number of native H-bonds (2D) |
| Folding target | Cα-RMSD:<br>1 Å (NTL9),<br>1.25 Å (Protein G)<br>Native H-bonds:<br>13 (NTL9),<br>10 (Protein G) |
| Bin Mapping | Rectilinear binmapper |

*Bin Mapping:*

<u>NTL9 (1D, Cα-RMSD, Å)</u>:
0.0 - 1.0 (1 bin), 1.0 - 4.4 (35 bins), 4.4 – 6.6 (12 bins), 6.6 – 10.2 (5 bins), 10.2 – inf (1 bin)

<u>NTL9 (2D, Cα-RMSD (Å) + Number of native H-bonds)</u>:
Cα-RMSD (Å): 0.0 - 1.0 (1 bin), 1.0 - 4.4 (35 bins), 4.4 – 6.6 (12 bins), 6.6 –10.2 (5 bins), 10.2 – inf (1 bin)
Number of native H-bonds: 0 – 9 (1 bin), 9 – 13 (5 bins), 13 – inf (1 bin)

<u>Protein G (2D, Cα-RMSD (Å) + Number of native H-bonds)</u>:
Cα-RMSD (Å): 0.00 – 1.25 (1 bin), 1.25 – 4.00 (36 bins), 4.00 – 7.00 (30 bins), 7.00 – 10.0 (15 bins), 10.0 – 13.0 (6 bins), 13.0 – inf (1 bin)
Number of native H-bonds: 0 – 5 (1 bin), 5 – 10 (5 bins), 10 – inf (1 bin)

*Computational Resources:*

NTL9 (γ=5ps$^{-1}$; 10 independent WESTPA simulations on 1 GPU each):
~22 days per WESTPA simulation
~24 minutes per iteration
Total number of iterations per WESTPA simulation: 1,200
Possible number of trajectories per iteration: 1456 (4 trajectories/bin x 364 possible 2D-bins)
Total number of trajectories over all iterations: ~1,200,000 (average WESTPA simulation)

NTL9 (γ=80ps$^{-1}$; 30 independent WESTPA simulations on 1 GPU or on 48 CPUs each):
~20 days per WESTPA simulation
~7 minutes per iteration
Total number of iterations per WESTPA simulation: 4,500
Possible number of trajectories per iteration: 210 (4 trajectories/bin x 52 possible 1D-bins)
Total number of trajectories over all iterations: ~850,000 (average WESTPA simulation)

Protein G (15 independent WESTPA simulations on 1 GPU each):
~31 days per WESTPA simulation
~30 minutes per iteration
Total number of iterations per WESTPA simulation: 1,500
Possible number of trajectories per iteration: 2436 (4 trajectories/bin x 609 possible 2D-bins)
Total number of trajectories over all iterations: ~1,700,000 (average WESTPA simulation)

**Flux profile calculation**

The profile of the flux, $J$, along the Cα-RMSD progress coordinate was calculated from the WE simulation data in two steps. First, inter-bin fluxes are calculated according to $\langle J_{ij}\rangle = \langle w_{ij}\rangle / \tau$, with $w_{ij}$ the weight transitioning from bin $i$ to bin $j$ in a single iteration and $\tau$ the resampling time (10 ps). Angular brackets indicate time- and ensemble-averages. Second, the net flux at the $i$th bin along the Cα-RMSD reaction coordinate was calculated from the difference in directional fluxes crossing from larger Cα-RMSD to smaller Cα-RMSD, $J_i = \sum_{j>i,k\leq i}(J_{jk} - J_{kj})$, where positive flux is directed towards the folded state and larger indices corresponding to larger RMSD values.

**haMSM analysis**

History augmented Markov State Models (haMSMs) were constructed directly from the WE simulation trajectories. Each haMSM was clustered using the latest-occurring 100,000 structures from the training period (Table S2). K-means clustering with a minimum-RMSD metric and all protein-atom Cartesian coordinates (rotationally and translationally minimized) was performed using the PyEMMA software package.[16] While haMSMs trained from steady-state trajectories yield mean first-passage times which are independent of the clustering and number of microstates,[10, 17] we found haMSMs with ~10,000 microstates were able to estimate steady-state kinetics from training data in the transient regime.[18] Given a set number of desired microstates and an initial condition, a k-means clustering is deterministic, but is random given a change in the desired number of microstates or initial bin. To quantify uncertainty in the clustering process, we therefore used three independent clusterings based on a set of similar numbers of microstates – i.e., 9999, 10000, and 10001.

Construction of the haMSM transition matrix proceeds in a highly similar manner to constructing a standard MSM, except via WE weights instead of simple transition counts. The mean first-passage times were obtained from the steady state flux of the haMSM transition matrix.[17] The lag time of the stochastic matrix was the same as the WE integration time, or 10.0 ps in this case, enabling the use of all of the stored trajectory information from the WE simulation to construct the haMSM. Further details can be found elsewhere.[18]

Error in WE-haMSM rate estimates was estimated in a manner analogous to the procedure for direct WE rate estimates (below), using a Bayesian bootstrapping approach. The haMSM flux matrix (transition matrix before row normalization) was calculated separately for each WE simulation, and each of the 3 independent clusterings. Bayesian bootstrapping (described below) was then performed by randomly weighting and summing together the flux matrix from each run, and was then row normalized to construct the bootstrap instance haMSM transition matrix and estimate the rate. For each clustering, 250 such Bayesian bootstrap samples were constructed, and the resulting 750 rates for each system were used to estimate 95% confidence intervals. This bootstrap ensemble contained 250 samples capturing the inter-run variability for each clustering, but only 3 clusterings to capture the variability of the clustering algorithm. As shown in Table S2, column 4, when equally weighting each WE run, the variability in the estimated rate between the clusterings is much less than the total error; thus, we do not expect additional clustering calculations to significantly change the measured confidence interval. The haMSM training period, estimated rate and confidence interval, and min/max estimated rate between the 3 clustering calculations with equally weighted transition information between each run, are shown in Table S2.

Table S2: haMSM analysis results

| System | Training period | haMSM folding MFPT | MFPT min/max from clustering |
|---|---|---|---|
| NTL9, $\gamma = 5\ ps^{-1}$ | 10-12 ns | 2.1 µs [0.78-9.0 µs] | 1.2-1.7 µs |
| NTL9, $\gamma = 80\ ps^{-1}$ | 30-45 ns | 0.44 ms [0.17-1.9ms] | 0.21-0.34 ms |
| Protein G, $\gamma = 5\ ps^{-1}$ | 10-15 ns | 0.014 s [.0033-.20s] | 0.016-0.027 s |

**Statistical analysis**

To assess the uncertainty of a WE-direct folding rate, a Bayesian credibility region[19] was calculated at each simulation time according to the Bayesian bootstrap procedure outlined by Rubin[20] and detailed by Mostofian et al.[21] The Bayesian bootstrap calculates the likelihood for a set of random parameters to describe the observed data based on a multinomial distribution and a uniform prior. For each set of rates, a 1000-fold bootstrap resampling was performed over multinomial parameters and the credibility region is reported as the interval that covered 95% of the weighted distribution of average rates. The Bayesian bootstrap is preferred in this study over the standard bootstrap method[22] which yields lower confidence limits that are systematically and significantly low.[21] However, it must be noted, that although we derive a nominal 95% credibility region for the observed data, this uncertainty interval, regardless of the procedure used, exhibits ~30% underestimation error and slight overestimation error which is intrinsic to the relatively small sample sizes.[21] Based on synthetic data resampled from the original data sets,[21] we conclude that the nominal 95% credibility region should be treated as a ~60% range of uncertainty. A larger range is not possible with the given small sample sizes. We emphasize, nevertheless, that possessing >10 truly independent and identically distributed values for a key observable is rather unusual and reflects the methodology choice and resource investment.

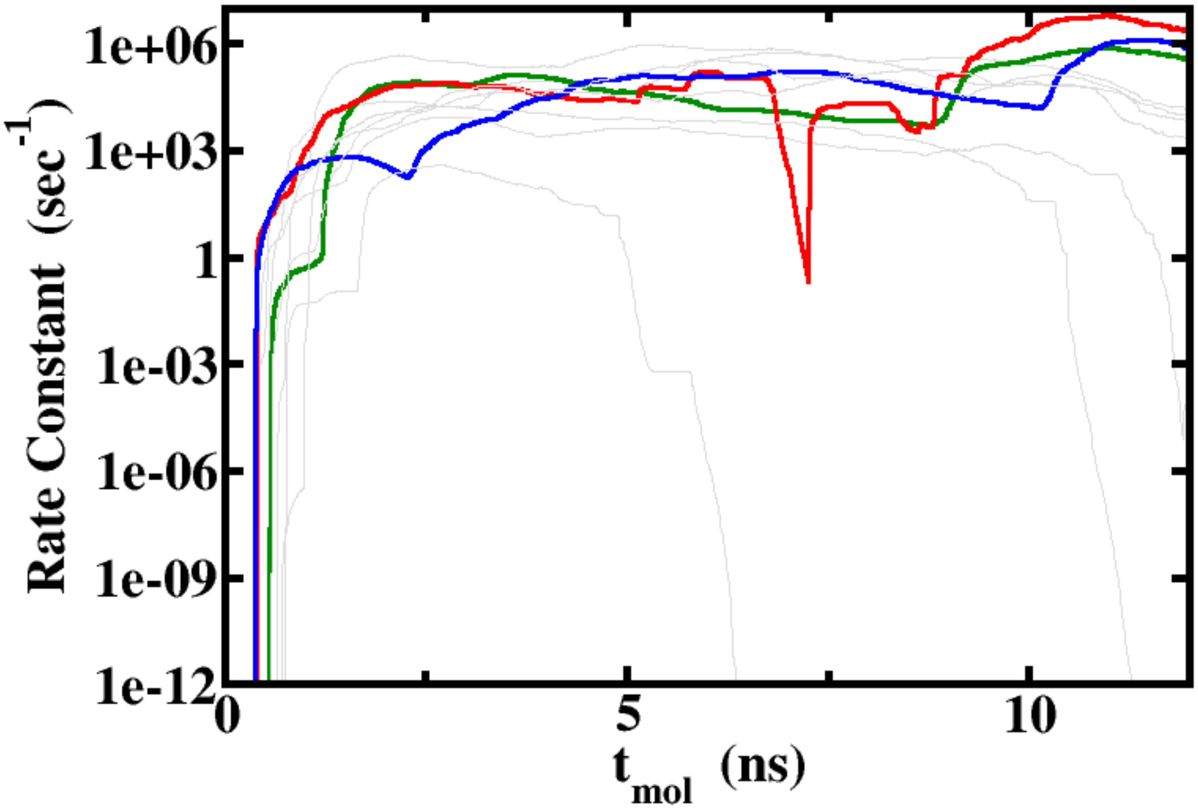


Figure S1: Fluctuations among WE runs. Evolution of the rate constant with molecular time for NTL9 folding using 2D WE method with solvent viscosity (γ) set to 5 $ps^{-1}$. Rate constants were calculated as windowed averages of the previous 1 ns. Color lines represent runs with top three highest rate constants and all other simulations are shown in gray.

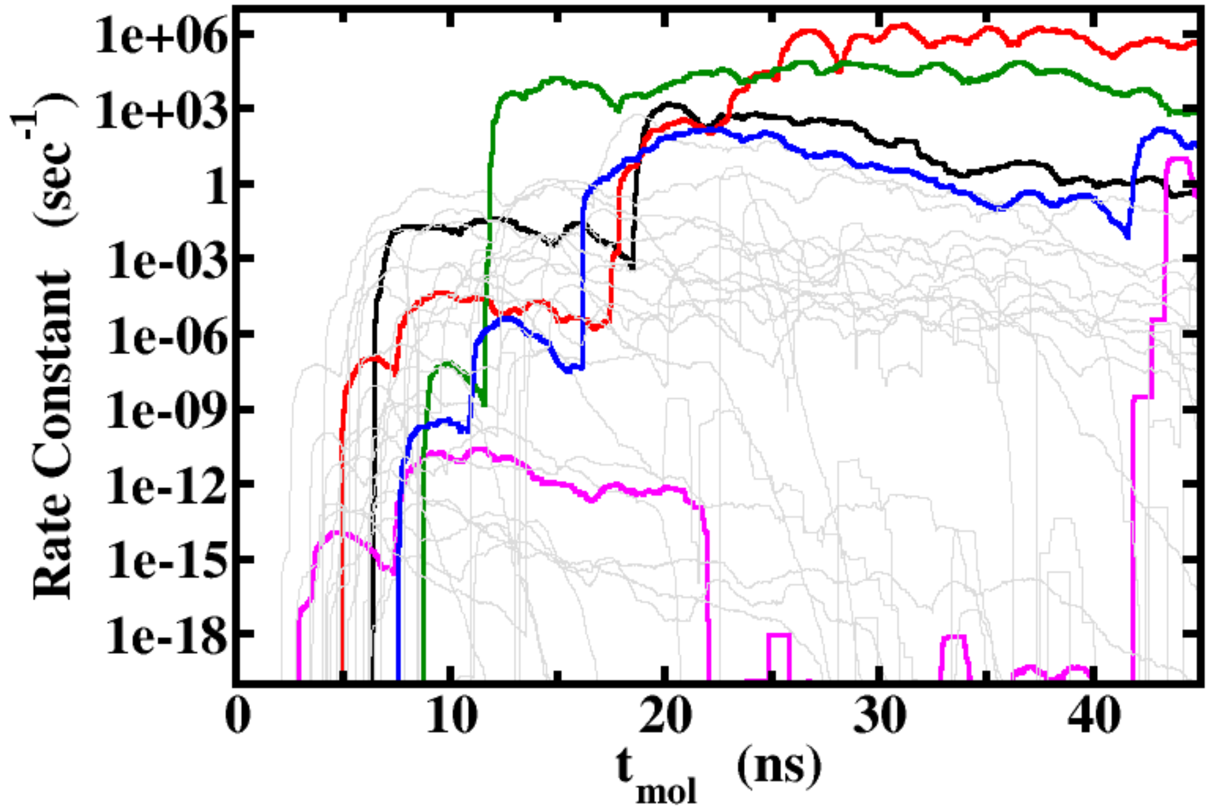


Figure S2: Fluctuations among WE runs. Evolution of the rate constant with molecular time for NTL9 folding using 1D WE method with solvent viscosity (γ) set to 80 $ps^{-1}$. Rate constants were calculated as windowed averages of the previous 1 ns. Color lines represent runs with top five highest rate constants and all other simulations are shown in gray.

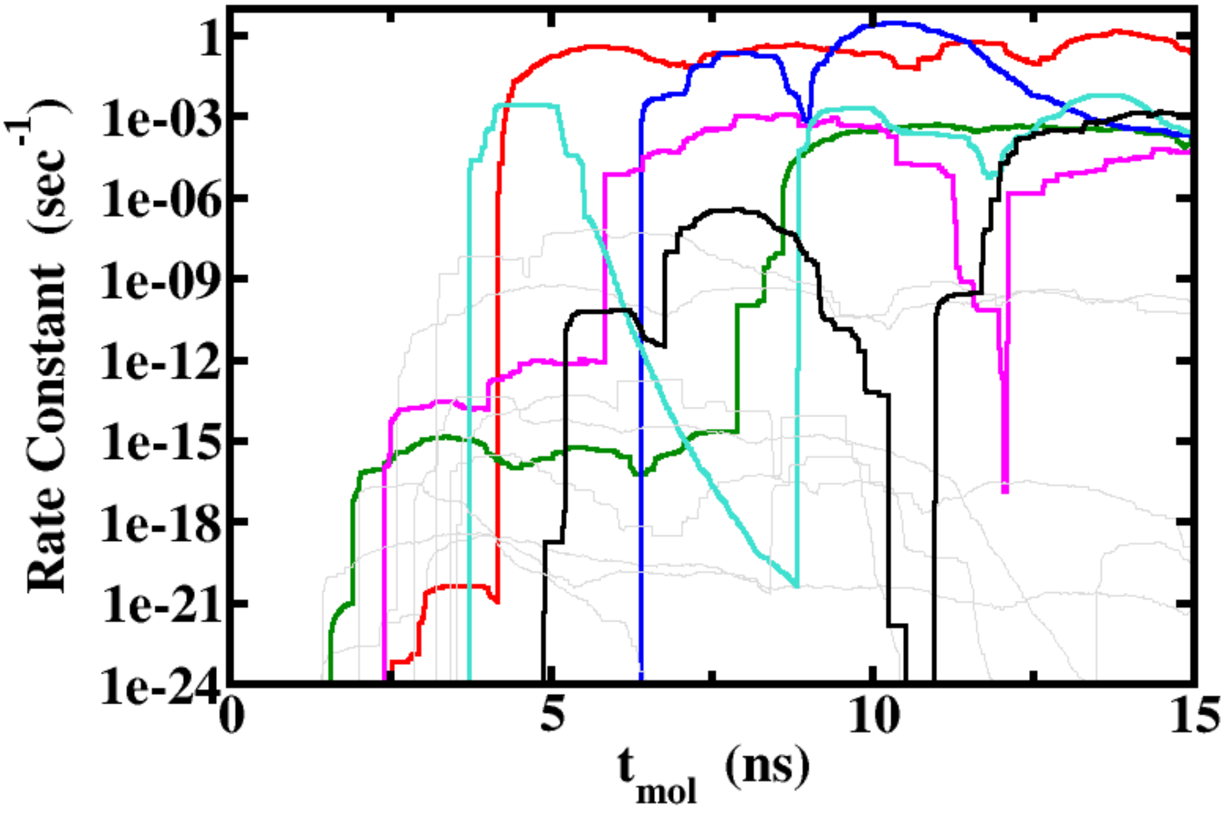


Figure S3: Fluctuations among WE runs. Evolution of the rate constant with molecular time for Protein G folding using 2D WE method with solvent viscosity (γ) set to 5 $ps^{-1}$. Rate constants were calculated as windowed averages of the previous 1 ns. Color lines represent runs with top six highest rate constants and all other simulations are shown in gray.

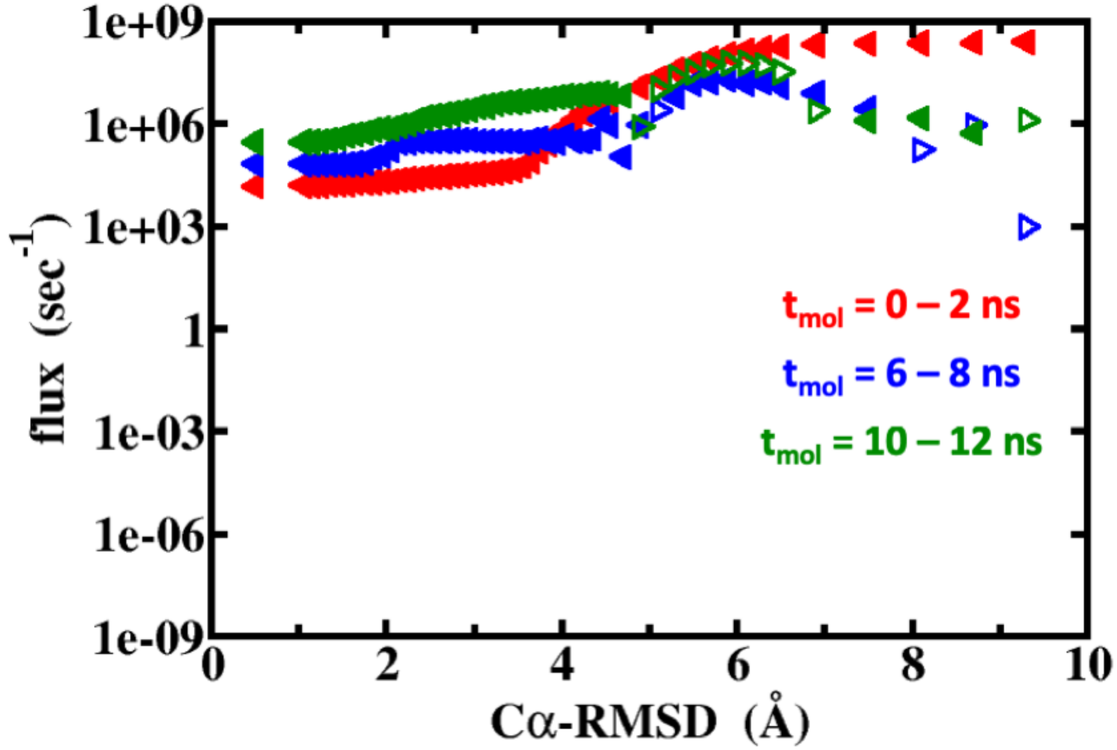


Figure S4: Flux profiles as a function of Cα-RMSD for NTL9 folding using 2D WE method, with solvent viscosity (γ) 5 $ps^{-1}$. The net change of probability, or flux, across a certain Cα-RMSD value during the given time range averaged over all 10 independent simulations is plotted as a function of that Cα-RMSD value. The filled left-pointing triangles indicate a flux toward the folded state and the empty right-pointing triangles indicate a flux toward the unfolded state. The “backwards” (unfolded-directed) flux values may reflect both noise in the data and that steady state has not been fully reached. We note that the flux profile is a fairly stringent assessment of the steady state that has not previously been used to assess WE simulations to our knowledge.

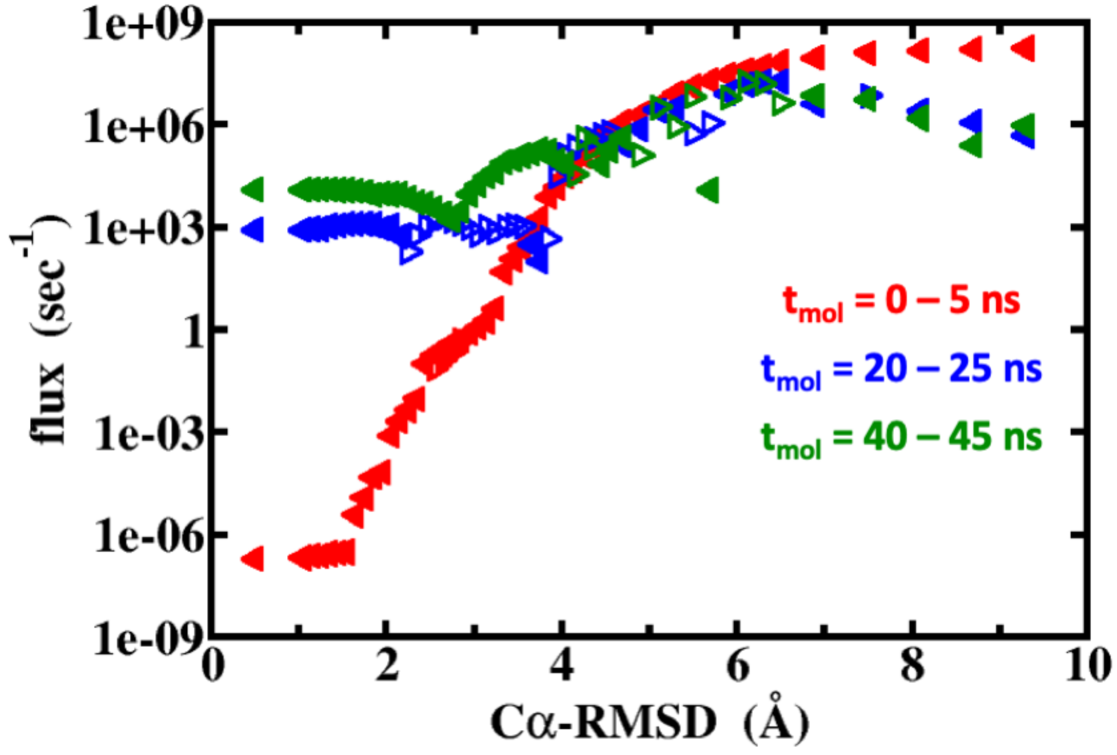


Figure S5: Flux profiles as a function of Cα-RMSD for NTL9 folding using 1D WE method, with solvent viscosity (γ) 80 $ps^{-1}$. The net change of probability, or flux, across a certain Cα-RMSD value during the given time range averaged over all 30 independent simulations is plotted as a function of that Cα-RMSD value. The filled left-pointing triangles indicate a flux toward the folded state and the empty right-pointing triangles indicate a flux toward the unfolded state. The "backwards" (unfolded-directed) flux values may reflect both noise in the data and that steady state has not been fully reached. We note that the flux profile is a fairly stringent assessment of the steady state that has not previously been used to assess WE simulations to our knowledge

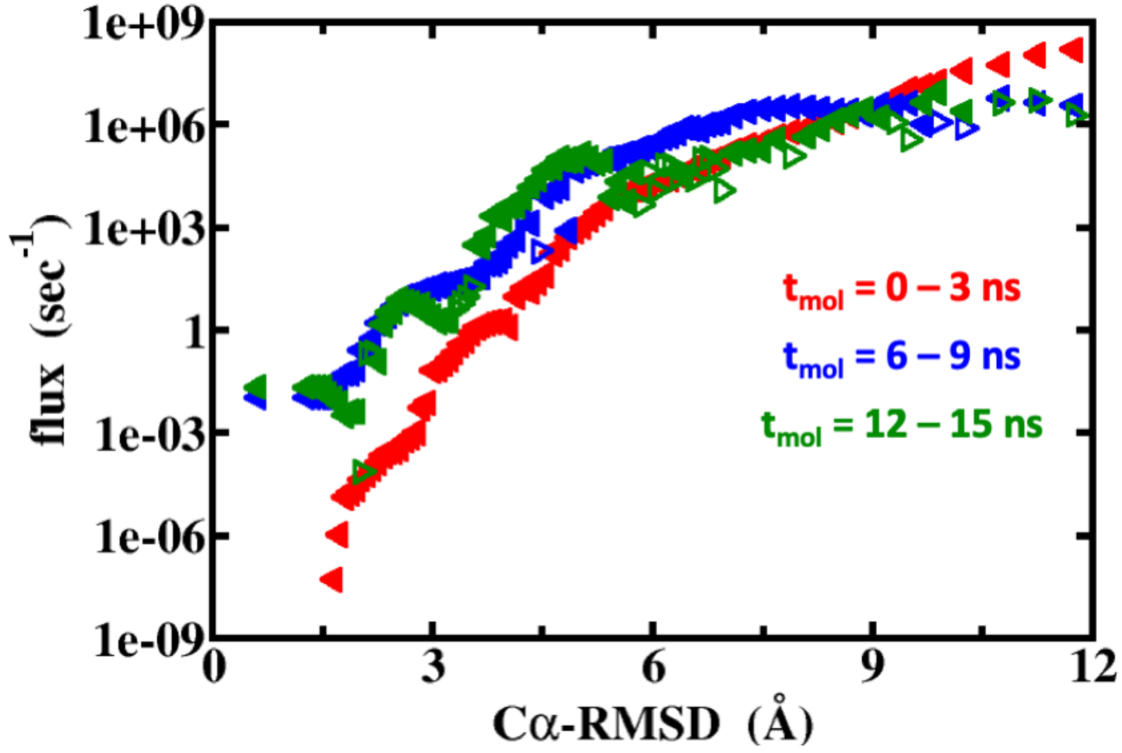


Figure S6: Flux profiles as a function of Cα-RMSD for Protein G folding using 2D WE method, with solvent viscosity (γ) 5 $ps^{-1}$. The net change of probability, or flux, across a certain Cα-RMSD value during the given time range averaged over all 15 independent simulations is plotted as a function of that Cα-RMSD value. The filled left-pointing triangles indicate a flux toward the folded state and the empty right-pointing triangles indicate a flux toward the unfolded state. The "backwards" (unfolded-directed) flux values may reflect both noise in the data and that steady state has not been fully reached. We note that the flux profile is a fairly stringent assessment of the steady state that has not previously been used to assess WE simulations to our knowledge

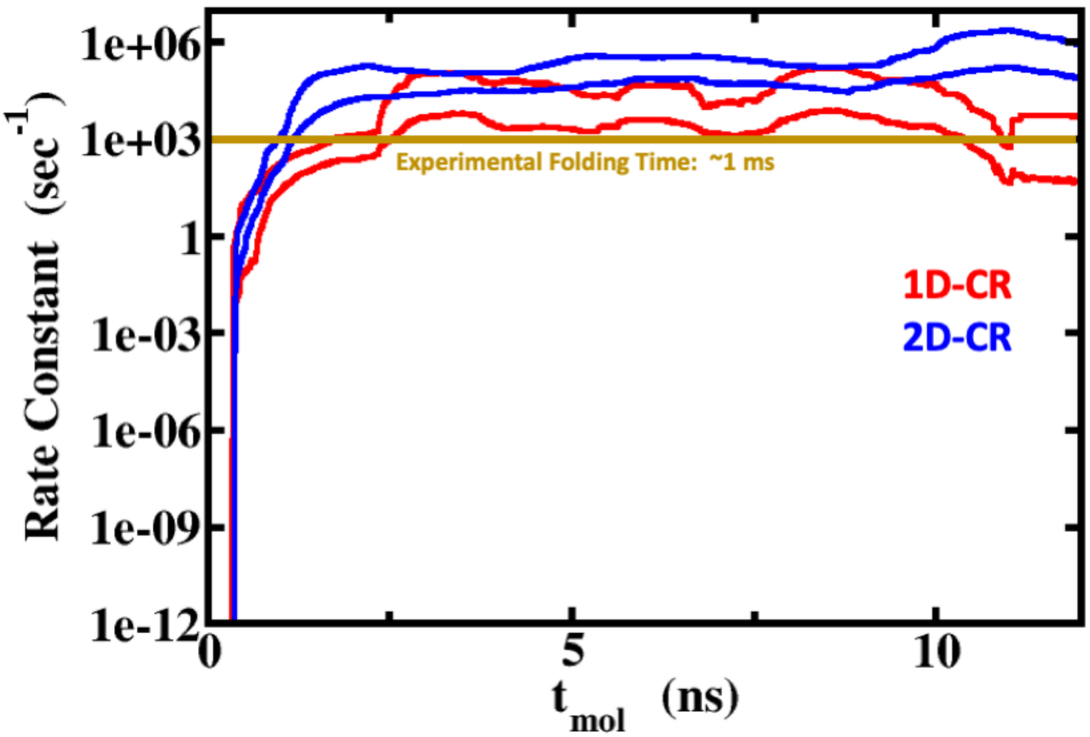


Figure S7: Comparison of two WE protocols for low-friction NTL9. Evolution of the average rate constant with molecular time for NTL9 folding with solvent viscosity (γ) set to 5 $ps^{-1}$ (aggregate time = 45 μs (1D, 30 independent simulations), 100 μs (2D, 10 independent simulations)). Rate constants were calculated as windowed averages of the previous 1 ns. The red (1D) and blue (2D) lines show the 95 % (nominal) Credibility Region (CR) calculated using the Bayesian bootstrap. Based on statistical re-analysis of the rate constants at $t_{mol}$ = 12 ns,[21] the Bayesian bootstrap tends to overestimate the true mean by ~5-8% and underestimates by ~36-39%.

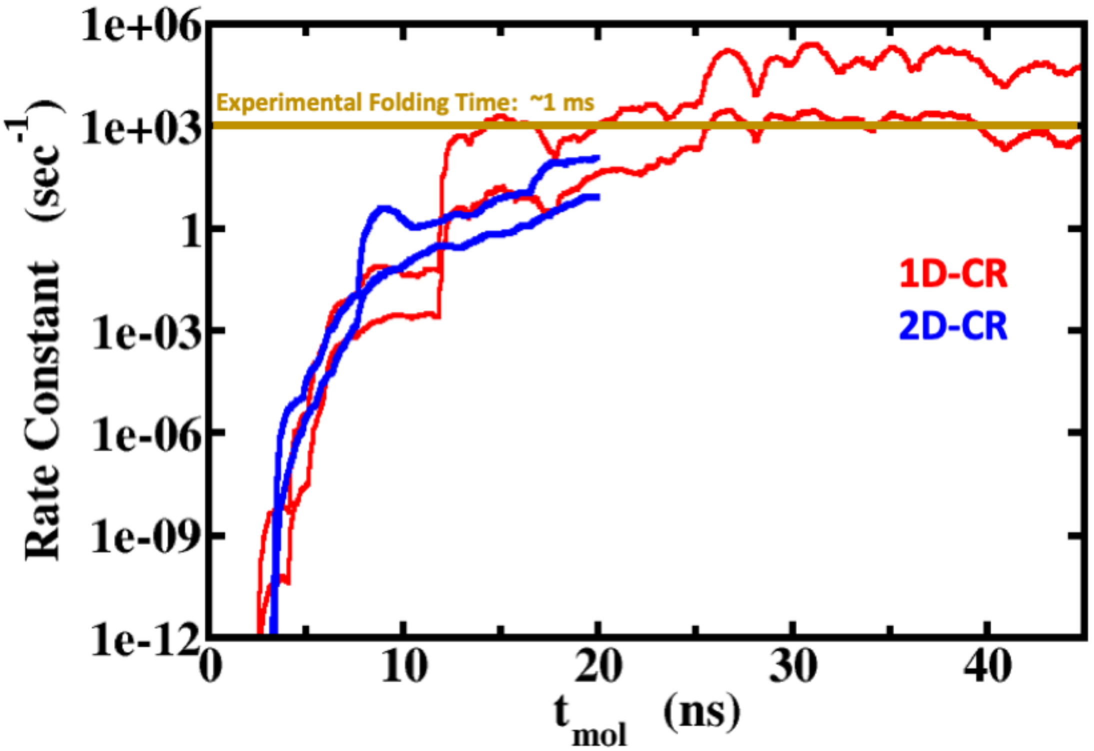


Figure S8: Comparison of two WE protocols for high-friction NTL9. Evolution of the rate constant with molecular time for NTL9 folding with solvent viscosity (γ) set to 80 $ps^{-1}$ (aggregate time = 252 µs (1D, 30 independent simulations), 200 µs (2D, 10 independent simulations)). Rate constants were calculated as windowed averages of the previous 1 ns. The red (1D) and blue (2D) lines show the 95% (nominal) Credibility Region (CR) calculated using the Bayesian bootstrap. Based on statistical re-analysis of the rate constants at $t_{mol}$ = 45 ns,[21] the Bayesian bootstrap overestimates the true mean by ~7-8% and underestimates by ~32-36% for 1D and 2D WE simulations, respectively.

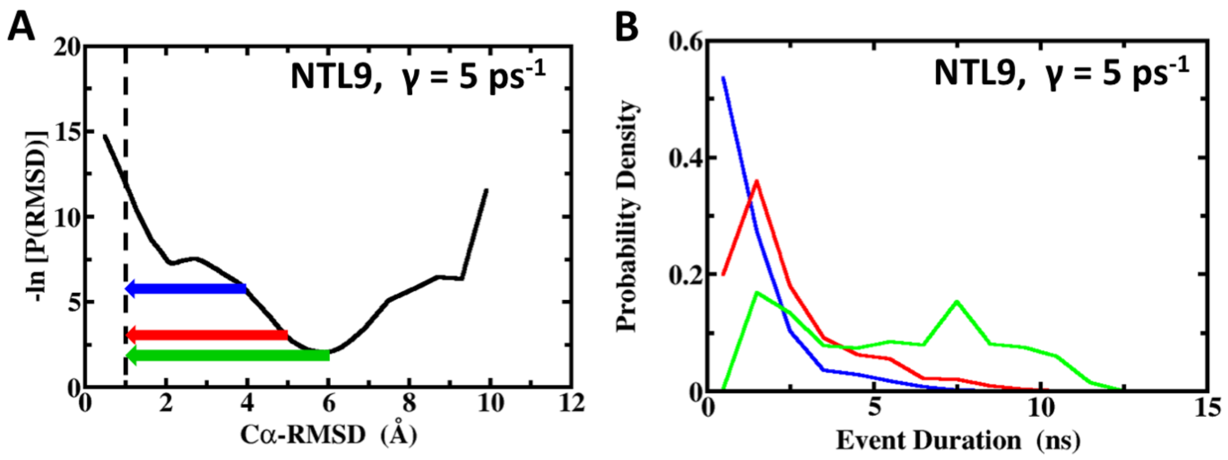


Figure S9: NTL9 folding events ($\gamma$ = 5 $ps^{-1}$): Definitions and durations. (A) Folding events are defined based on Cα-RMSD thresholds, and horizontal arrows indicate three choices examined for event definitions: starting at 4 Å (blue), 5 Å (red), and 6 Å (green). For reference, the black curve shows the negative logarithm of the weighted Cα-RMSD distribution, which serves as an effective non-equilibrium free energy profile of the steady-state folding simulation. Because of the absorbing boundary at Cα-RMSD = 1 Å, probability is extensively depleted from low Cα-RMSD values as compared to equilibrium, thus elevating the effective free energy. (B) An event duration is defined as the time span between the time point at which a structure had assumed the event starting point, i.e. a Cα-RMSD value of at least 4 Å (blue), 5 Å (red), or 6 Å (green), for the last time and the time point at which it reached the target state in that same continuous trajectory, weighted by the probability of the simulation when reaching the target state. The distributions, with a bin width of 1 ns, are based on ~80,000 events.

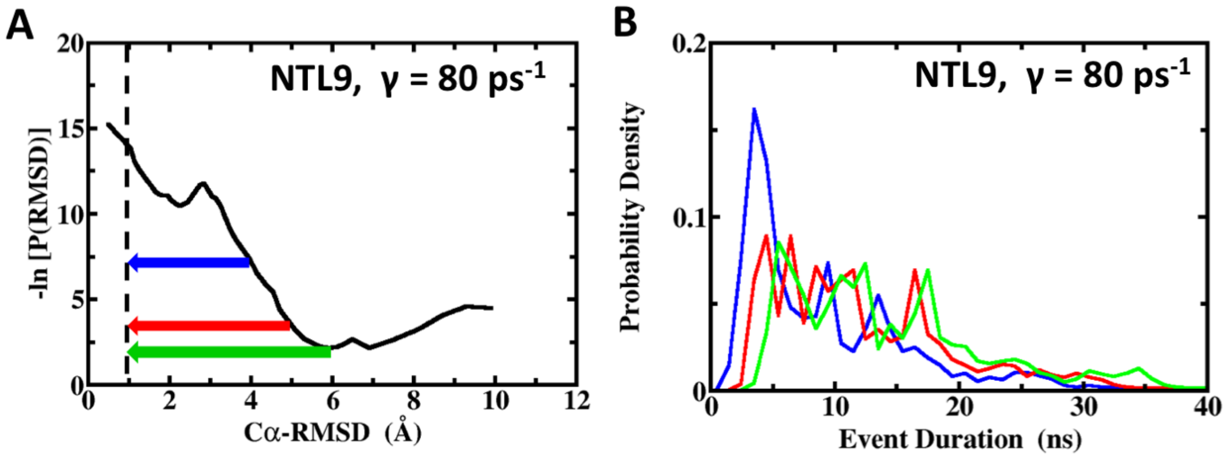


Figure S10: NTL9 folding events ($\gamma$ = 80 $ps^{-1}$): Definitions and durations. (A) Folding events are defined based on Cα-RMSD thresholds, and horizontal arrows indicate three choices examined for event definitions: starting at 4 Å (blue), 5 Å (red), and 6 Å (green). For reference, the black curve shows the negative logarithm of the weighted Cα-RMSD distribution, which serves as an effective non-equilibrium free energy profile of the steady-state folding simulation. Because of the absorbing boundary at Cα-RMSD = 1 Å, probability is extensively depleted from low Cα-RMSD values as compared to equilibrium, thus elevating the effective free energy. (B) An event duration is defined as the time span between the time point at which a structure had assumed the event starting point, i.e. a Cα-RMSD value of at least 4 Å (blue), 5 Å (red), or 6 Å (green), for the last time and the time point at which it reached the target state in that same continuous trajectory, weighted by the probability of the simulation when reaching the target state. The distributions, with a bin width of 1 ns, are based on ~65,000 events.

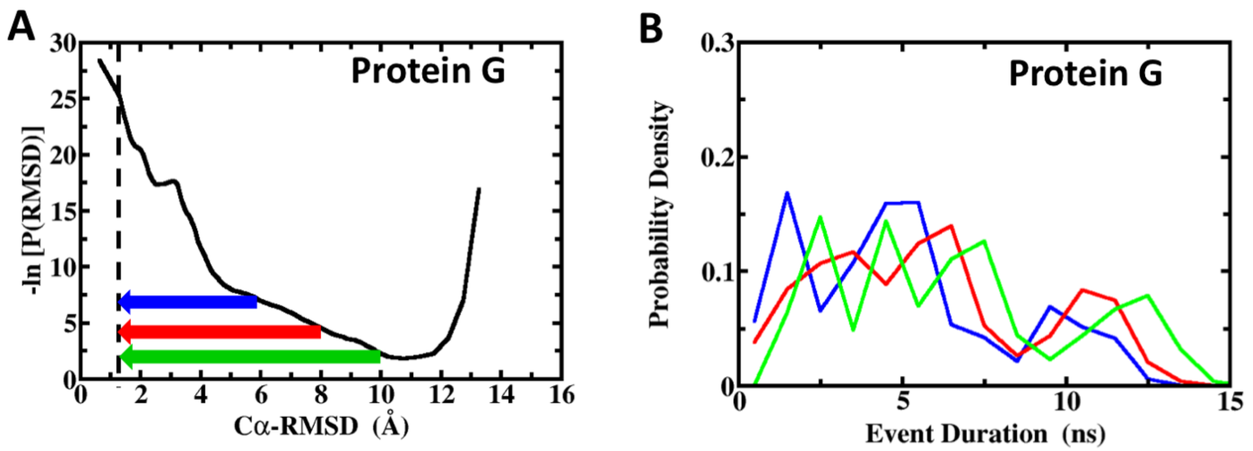


Figure S11: Protein G folding events: Definitions and durations. (A) Folding events are defined based on Cα-RMSD thresholds, and horizontal arrows indicate three choices examined for event definitions: starting at 6 Å (blue), 8 Å (red), and 10 Å (green). For reference, the black curve shows the negative logarithm of the weighted Cα-RMSD distribution, which serves as an effective non-equilibrium free energy profile of the steady-state folding simulation. Because of the absorbing boundary at Cα-RMSD = 1.25 Å, probability is extensively depleted from low Cα-RMSD values, as compared to equilibrium, thus elevating the effective free energy. (B) An event duration is defined as the time span between the time point at which a structure had assumed the event starting point, i.e. a Cα-RMSD value of at least 6 Å (blue), 8 Å (red), or 10 Å (green), for the last time and the time point at which it reached the target state in that same continuous trajectory, weighted by the probability of the simulation when reaching the target state. The distributions, with a bin width of 1 ns, are based on ~50,000 events.

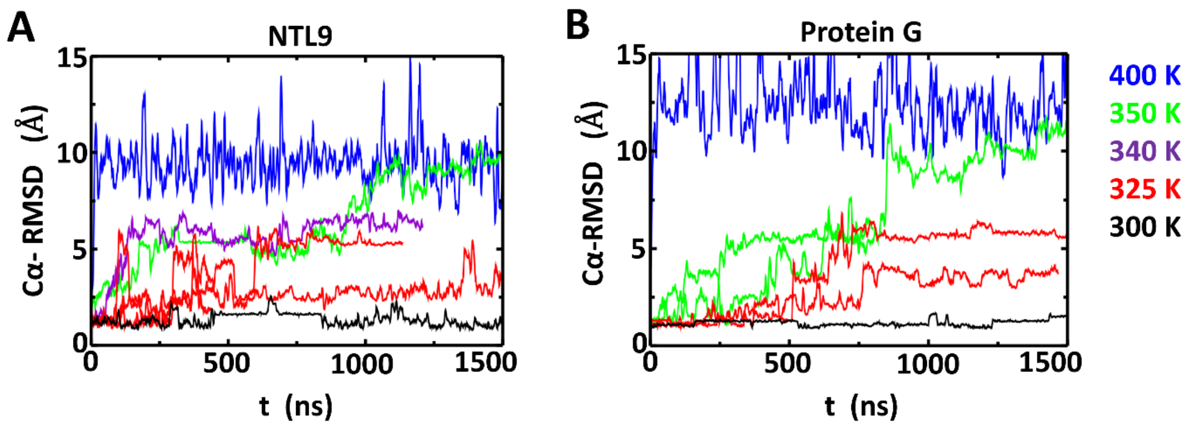


Figure S12: Estimation of approximate melting/folding temperatures. Plotted is the time evolution of Cα-RMSD from brute-force simulations of (A) NTL9 and (B) Protein G at temperatures ≥ 300 K. The simulations were at least 1 μs long to assess the stability of the proteins at the corresponding temperature. Repeated simulations show that both proteins can more frequently unfold and re-fold at 325 K compared to other temperatures. Thus, we roughly estimate Tm ~ 325 K to be the melting temperature of both proteins, which is somewhat lower than the experimental results (355 K for NTL9[23] and 365 K for Protein G[24]).

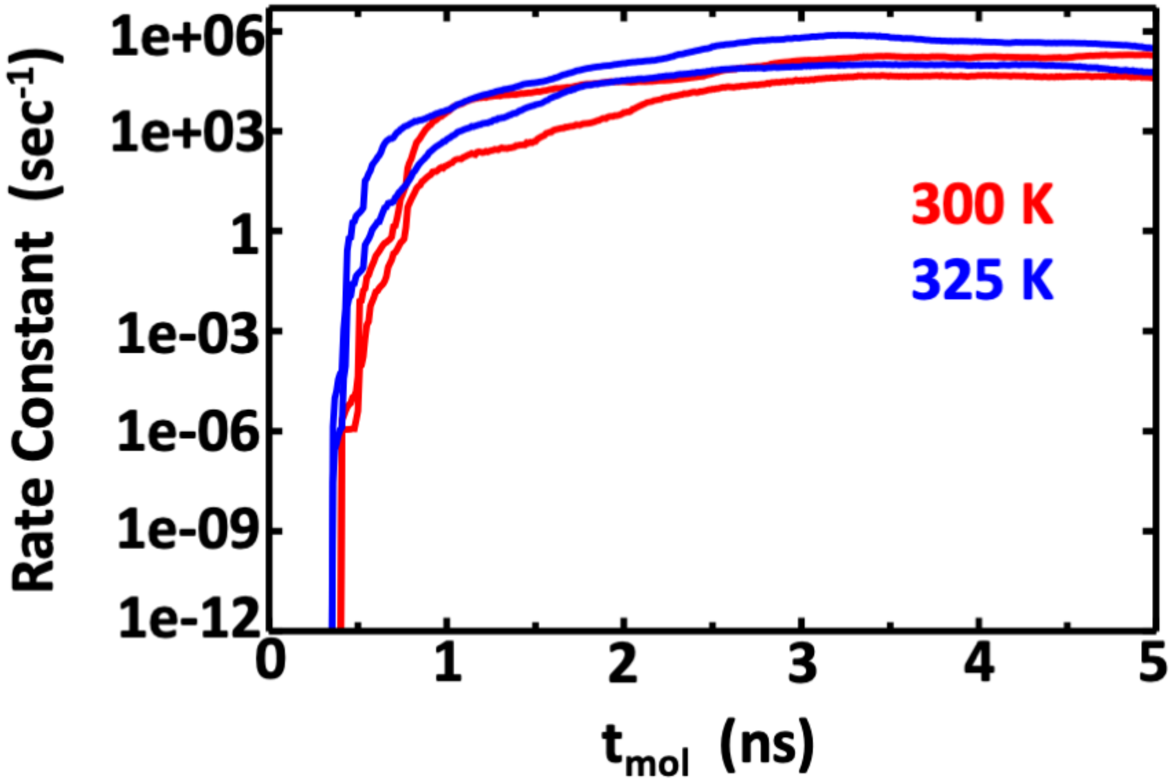


Figure S13: Rate constant estimations for NTL9 folding at 300 K (red) and at 325 K (blue) using 2D WE method with solvent viscosity (γ) set to 5 ps$^{-1}$. The lines show the nominal 95% Credibility Region (CR) as a function of molecular time from Bayesian bootstrapping based on direct WE rate constant estimates, which were windowed averages of the previous 1 ns of molecular time for each of the 5 independent simulations. The credibility region is shifted to slightly larger rate constants at 325 K compared to 300 K, yet the two profiles are overlapping for the most part. This suggests that NTL9 does not fold significantly faster at its melting temperature than it does at room temperature.
